# Patterns and drivers of pollen co-transport networks vary across pollinator groups

**DOI:** 10.1101/2023.09.20.558666

**Authors:** Liedson Tavares Carneiro, Jessica Nicole Williams, Daniel A Barker, Joseph W Anderson, Carlos Martel, Gerardo Arceo-Gomez

## Abstract

Pollen transport is an understudied process with consequences for plant reproductive success and floral evolution. Recently, pollinator bodies have been described as pollen competition arenas, with implications for plant community assembly. However, the identity, strength, and diversity of pollen competitive interactions and how they vary across pollinator groups is unknown. Evaluating patterns and drivers of the pollen competition landscape across different pollinator groups is central to further our understanding of plant coexistence mechanisms. Here, we integrate information on insect pollen loads with network analyses to uncover novel pollen co-transport networks and how these vary across pollinators. We evaluate differences in pollen load size, diversity and their phenological and phylogenetic attributes among insect groups and how these relate to body size and sex. Pollen co-transport networks revealed differences in the number and identity of competitors that pollen species encounter across pollinator groups. These networks were highly modular, with groups of pollen species interacting more often on pollinator bodies. Pollen load size and richness were shaped by bee size and sex. Sex also influenced the pollen phylogenetical diversity. Pollinators can impose vastly different competitive landscapes during pollen transport, with so far unknown consequences for plant reproductive success, floral evolution and community assembly.

## 1 Introduction

The pollination process has been the subject of ample study for over 200 years (Darwin, 1877; Mayr, 1986; Vogel, 1996; Waser & Ollerton, 2006). As a result, much is known about the factors that influence pollen removal from anthers (*e.g*., pollen production and presentation), its deposition on stigmas (*e.g*., pollinator attraction and efficiency) (Harder & Routley, 2006; Morales & Traveset, 2008; Armbruster *et al*., 2009; Moreira-Hernández & Muchhala, 2019), and pollen germination and ovule fertilization (*e.g*., stigma receptivity and pollen competition) (Dafni & Maués, 1998; Arceo-Gómez & Ashman, 2011; Streher *et al*., 2020; Lopes *et al*., 2022). Together, these represent the start and endpoints in the pollination process and have direct implications for pollination success and floral evolution. However, less is known about more intermediate stages in the pollination process, such as those that inform on the drivers, patterns, and consequences of pollen transport on insect bodies (but see Minnaar *et al*., 2019 and references therein). The study of pollen transport alone can help uncover novel mechanisms affecting male and female success in plants (Minnaar *et al*., 2019) and pollinator foraging niche breadth (*e.g*., Smith *et al*., 2019). The study of pollen transport patterns can also shed light into the evolution of pollen dispersal strategies (Minnaar *et al*., 2019), or provide evidence of community-level direct and indirect plant–plant interactions (*e.g*., competition) (*e.g*., Lázaro et al. 2014, Arceo□Gómez et al. 2016, Cullen et al. 2021). More recently, insect bodies have been described as ‘competitive arenas’, where co-transported pollen grains can compete for space (Minnaar *et al*., 2019; Moir & Anderson, 2023). Nevertheless, the diversity, strength, and identity of pollen–pollen interactions on insects, and how it varies across insect taxa, is not well-known. Evaluating the pollen ‘landscape’ (*e.g*., pollen load size and diversity) on insect bodies, as well as the ecological and phylogenetic attributes of pollen loads carried across insect groups, has the potential to further advance our understanding of coexistence mechanisms and drivers of plant community assembly.

Knowledge of the size, diversity and identity of pollen loads carried by flower-visiting insects can be used to inform how pollinators may differentially mediate ecological and evolutionary processes within communities. Patterns of pollen transport (load size and diversity) and co-transport (identity and frequency of co-transported pollen grains) may vary greatly among insect species or taxonomic groups (Zhao *et al*., 2019) depending on their morphology, nutritional needs, foraging behavior and life history (Alarcon, 2010; Vaudo *et al*., 2020; Cullen *et al*., 2021). Bees, for example, are considered more efficient at pollen transport compared to other non-bee flower-visiting insects due to their strong dependence on nectar and pollen (Ollerton, 2017; Zhao *et al*., 2019). Flies, on the other hand, generally exhibit lower pollen transport efficiency (Alarcon, 2010; Zhao *et al*., 2019; Cullen *et al*., 2021). Although, studies evaluating the importance of flies as pollinators are limited (Földesi *et al*., 2021). Even within bees, a wide range of morphologies, foraging behaviors and resource preferences likely lead to differences in patterns of pollen transport. For instance, eusocial bees such as honeybees and bumblebees are known to have wide niche breadths (Kleinert & Giannini, 2012; Johnson & Ashman, 2019; Wood *et al*., 2021) likely leading to the transport of large and diverse pollen loads. If this is the case, however, the body of eusocial bees may also represent a harsher competitive arena for pollen grains compared to that of solitary bees with narrower foraging niches (Grüter & Hayes, 2022), with potential consequences for the evolution of pollen dispersal strategies in plants (Minnaar *et al*., 2019).

Furthermore, while insect species can be generalist and carry diverse pollen loads, individual insect loads may be composed of only one or a few pollen species (Cane & Sipes, 2006; Leonhardt & Blüthgen, 2012; Smith *et al*., 2019). Foraging niche partitioning among individual insects within a species or taxonomic group may have the potential to generate differences in the identity and strength of interactions between pairs of pollen species; *i.e*., co-transport (Brosi, 2016; Smith *et al*., 2019). Evaluating patterns of pollen co-transport at the individual level is thus key for understanding how pollinator species can generate different pollen competition landscapes (*e.g*., Minnaar et al. 2019). Here, we integrate information on the number and identity of pollen grains on individual insect bodies with network analytical tools to reveal detailed pollen co-transport networks (Figure 1), and evaluate how their structure varies across pollinator taxa. This approach can also help identify groups of pollen species that travel, and likely compete with each other, more often than with other pollen species (*i.e*., modularity; Olesen *et al*., 2007) and how these pollen–pollen interactions change across different pollinators. Overall, this approach has the potential to help further our understanding of how different pollinator groups can impose different ‘pollen competitive arenas’. That is, how the identity, strength, and diversity of competitive interactions that pollen species encounter changes as they travel across different pollinator bodies, hence affecting subsequent stages in the pollen pathway (Minnaar *et al*., 2019). Such knowledge will help advance our understanding of patterns and drivers of pollen co-transport among individual insects and pollinator groups (Smith *et al*., 2019; Cullen *et al*., 2021), as well as its potential ecological and evolutionary consequences.

**Figure 1.**
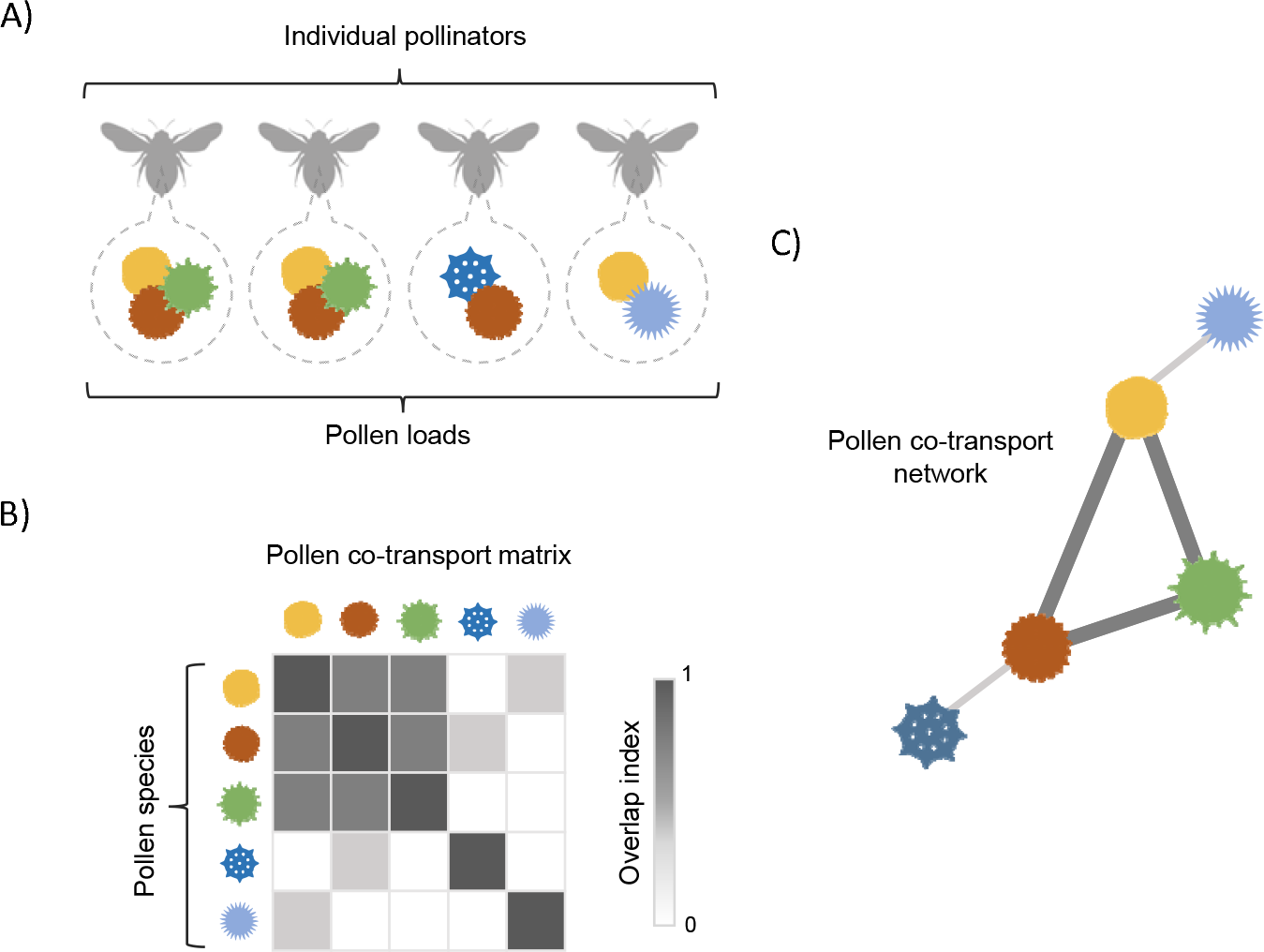
Conceptualization and construction of pollen co-transport networks. A) Pollen grains within pollen loads carried by individual insects within pollinator groups are identified and quantified. B) A pollen co-transport matrix is created using Schoener’s overlap index to quantify the strength of co-transport between each pollen species pair. C) A one-mode network is built with pollen species as nodes and overlap index values (co-transport strength) as weighted links.

Pollinator body size and sex can further help mediate differences in patterns of pollen transport among bees. Body size for instance can vary widely within and among bee species (Michener, 2007; Danforth, 2019) and is directly related to foraging distance as well as to their capacity to carry large pollen loads (Greenleaf *et al*., 2007; Wright *et al*., 2015). Body size has also been shown to influence the range of floral phenotypes that bees can access (Solís-Montero & Vallejo-Marín, 2017), hence influencing the number, diversity and composition of pollen loads. Sex-related differences in bee behavior and resource preferences also have the potential to play an important role in shaping patterns of pollen transport (Roswell *et al*., 2019; Cullen *et al*., 2021). Unlike eusocial male bees, male solitary bees not only forage for nectar, but also use flowers for patrolling, mating and sleeping (Pinheiro *et al*., 2017; Danforth, 2019), potentially increasing the diversity of pollen loads. Sex-based differences in pollen transport can further arise because females are active pollen collectors that contact anthers more often compared to males (Danforth, 2019). Evaluating the phenotypic drivers underlying variation in patterns of pollen transport is central for advancing our understanding of its potential ecological (Ne’eman *et al*., 2006; Cullen *et al*., 2021) and evolutionary consequences (Minnaar *et al*., 2019).

Beyond their taxonomic identity and abundance, pollen grains can provide information on the ecological attributes and evolutionary history of the plants transported by insects (Cullen *et al*., 2021; Wood *et al*., 2021), which can help assess the importance of different insect groups across temporal (Yourstone *et al*., 2023) and phylogenetic scales (Srivastava *et al*., 2012). For instance, an unexplored component of the ‘pollen landscape’ on insect bodies is the phenological diversity of pollen loads. Eusocial bees typically forage for longer times over the season and can thus be expected to carry pollen from species with large phenological differences (Danforth, 2019; Glaum *et al*., 2021; Yourstone *et al*., 2023). Pollinators with pollen loads that have a large phenological diversity (*i.e*., different flowering times) can be key in structuring and maintaining plant communities across temporal scales (Koski *et al*., 2015). Insects carrying phenologically diverse pollen loads, however, may also contribute to strengthen competitive plant–plant interactions via heterospecific pollen transfer. For instance, heterospecific pollen transfer effects can increase if pollinators facilitate transfer of pollen among plant species with little to no flowering overlap (*e.g*., Waser, 1978; Arceo-Gómez, 2021). Similarly, evaluating the phylogenetic diversity of pollen loads carried can help inform on key ecological and evolutionary process within communities. For instance, large phylogenetic diversity within pollen loads may suggest high evolutionary convergence across plant species to attract similar pollinators (Wood *et al*., 2021), and/or high niche differentiation among individual insects within a species or taxonomic group (Smith *et al*., 2019). Hence contributing to the maintenance of phylogenetically diverse plant communities (Vieira *et al*., 2013). From an ecological perspective, pollen loads composed of closely related plant species (*i.e*., low phylogenetic diversity) may lead to stronger plant–plant competition via heterospecific pollen transfer (Arceo-Gómez & Ashman, 2011; Arceo□Gómez *et al*., 2016), as its negative fitness effects increase with increasing phylogenetic relatedness (Arceo□Gómez & Ashman, 2016; Streher *et al*., 2020). Finally, high phylogenetic diversity in pollen loads may also indicate that insect bodies may act as competitive arenas for distantly related pollen grains, with unknown consequences for the evolution of pollen dispersal strategies. Evaluating the extent and variation in phenological and phylogenetic diversity of pollen loads carried across pollinator taxa is thus an important step in elucidating its potential ecological and evolutionary importance in co-flowering communities.

In this study, we evaluate whether phenotypic or life-history traits across different pollinator groups play a role in shaping patterns of pollen transport (load size and diversity) and co-transport within a diverse co-flowering community in Northern California. Particularly, we evaluated how pollen load size and richness, its phenological and phylogenetic assembly, as well as pollen co-transport network structure vary across different pollinator groups. We further evaluated whether differences in pollen load size, richness, phylogenetic and phenological diversity among bees are determined by body size and sex.

## 2 Materials and Methods

### 2.1 Study site and system

The study was carried out at the McLaughlin Natural Reserve, Lower Lake, CA, USA (38°51’41.4”N, 122°23’54.8”W, alt. = ∼650 m) during the peak flowering season of the serpentine seep metacommunity at two sites (BS and RHA; see Wei *et al*., 2021) in 2021. Studied sites combined comprise ∼50 herb and subshrub species growing within grasslands and shrublands (Koski *et al*., 2015), and ∼200 flower-visiting insect species (Carneiro *et al*. unpublished data; also see Wei *et al*., 2021). The flowering season in the seeps is short, occurring between May and July (Arceo-Gómez *et al*., 2018). Previous studies at these seeps had shown the high importance of direct and indirect pollinator mediated plant–plant interactions in mediating plant community assembly (*e.g*., Koski *et al*., 2015; Arceo□Gómez *et al*., 2016; Arceo-Gómez *et al*., 2018; Albor *et al*., 2020, 2022).

### 2.2 Pollinator and pollen load sampling

Flower-visiting bees and flies were collected foraging on the serpentine seeps between 09h00 and 15h00 using entomological nets. Collections took place each day by alternating sites over 13 days between May 9^th^ and June 1^st^, 2021. Insects were collected by 2-3 people simultaneously by walking at a steady pace while observing all plant species within each site and collecting every insect observed visiting a flower and making contact with the reproductive structures. Specimens were stored in tubes under freezing temperatures from collection to processing in the lab. Pollen loads were collected from insect bodies (head, proboscis, dorsal and ventral thorax, and fore- and mid-legs) using fuchsin jelly cubes that were later mounted on microscope slides (Beattie, 1971). Pollen carried on bee corbiculae or scopae was excluded because it represents a resource that is not directly available for pollination (Tong & Huang, 2018; Weinman *et al*., 2023). All pollen grains obtained from insects were counted under a microscope and identified based on pollen libraries previously established from anthers collected for each plant species at the study sites. We obtained pollen load size, richness and composition for each individual insect. Insects were separated into morphospecies and grouped into five major categories that represent their taxonomical identity as well as differences in their morphology and foraging behavior, all of which can impact pollen transport patterns (Smith *et al*., 2019; Cullen *et al*., 2021). The following pollinator groups were used: bumble bees, honeybees, megachilid bees, other bees (non-megachilid non-eusocial bees) and flies. Bee specimens were identified as female or male and taxonomically at the genus level using identification keys (Michener *et al*., 1994; Michener, 2007) prior to morphospecies assignment. We also measured bee intertegular distance (ITD) as a surrogate of body size (Cane, 1987). Insect specimens are preserved in the insect collection at the East Tennessee State University (ETSU).

### 2.3 Pollen co-transport network

To evaluate differences in the structure of pollen co-transport networks across pollinator groups, we restricted our analysis to insects that were collected at the most well-sampled site (RHA). For each pollinator group (except ‘other bee’ category), we used the number and identity of pollen grains on each individual insect collected to construct weighted pollen co-transport networks (Figure 1A-C). For this, we estimated the degree of ‘insect body use’ overlap (*i.e*., the strength of co-transport) (Figure 1B) and used it as a link between plant species whose pollen grains were observed on individual insects within each pollinator group (Figure 1C). The strength of co-transport was calculated for each pollen species pair as the Schoener’s niche overlap index (SI) (Schoener, 1970; Linton *et al*., 1981), which is widely used in ecological research (*e.g*., Forrest 2015, Arceo-Gómez et al. 2018, Albor et al. 2020, 2022). Schoener’s niche overlap index was estimated for each pollen species-pair and within each pollinator group. A higher degree of co-transport indicates a higher degree of sharing of insect bodies between two pollen species within a given pollinator group. We constructed one-mode networks using the *qgraph* R package (Epskamp *et al*., 2012) and the co-transport values (Schoener’s niche overlap index) as weighted links between pollen species. For each pollen co-transport network, we estimated pollen co-transport degree (*i.e*., total number of co-transport partners) and strength (*i.e*., intensity of ‘body use’ overlap with other pollen species) for each pollen species. We also estimated modularity (*Q*) to uncover groups of pollen species that travel more often with each other on insect bodies than with other species within each pollen co-transport network using the optimal community structure algorithm through modularity maximization (Brandes *et al*., 2008).

### 2.4 Phenological diversity

We estimated flowering time distances between all plant species pairs found at the study sites. For this, we quantified the production of flowers per species over a two-week sampling period in the flowering season of 2021. We set 1×2-meter plots along the seeps totalizing 21 plots at BS site and 19 plots at RHA site. Plant identity and number of open flowers per species were recorded within plots every sampling day. The number of open flowers per plant species per day was used to calculate the Schoener’s niche overlap (SI) between each species pair representing the temporal overlap between species as in previous studies (e.g., Arceo-Gómez *et al*., 2018; Albor *et al*., 2020, 2022). The temporal overlap values were transformed to distance values (1 □ SI) representing flowering time distances between plant species. For insect pollen loads that contained three or more plant species (*n* = 359), we calculated mean flowering time distance (MFTD) which ranged from 0 to 1. Larger values represent greater phenological distance among plant species sharing an insect body, hence indicating greater phenological diversity.

### 2.5 Phylogenetic diversity

We built a phylogenetic tree for each insect pollen load containing three or more plant species using the function *phylo.maker* in the R package *V.PhyloMaker* (Jin & Qian, 2019), specifying the mega-tree of vascular plants ‘GBOTB.extended’ as the source tree (Smith & Brown, 2018). To estimate phylogenetic diversity of pollen loads transported by insects we obtained pairwise phylogenetic distances between plant species composing each pollen load using the function *cophenetic*. From each insect pollen load we computed the mean phylogenetic distance (MPD) adjusted by the proportional product of the abundances (*i.e*., number of pollen grains) of plant species presented in the matrix using the function *mpd* from the R package *picante* (Webb *et al*., 2002; Kembel *et al*., 2010).

### 2.6 Data analyses

We modeled pollen load size, richness, MPD and MFTD as a function of pollinator category, sex and body size using linear mixed models. For pollen load size and richness, we used generalized linear mixed models (GLMM) for count data. Since models accounting for Poisson error-distribution were overdispersed, we fitted a negative binomial GLMM using *glmmTMB* (R package glmmTMB, Brooks *et al*., 2017), which improved all model parameters and fit. When modelling pollen load size and richness as a function of pollinator body size, we also considered its interaction with pollinator group to evaluate whether body size effects depend on pollinator group. For weighted MPD and MFTD, we used Gaussian models using the *lmer* function (R package lme4, Bates *et al*., 2015). The variation clustered by insect samples belonging to the same morphospecies was considered by adding morphospecies identity as a random effect in all models. To test for differences in the same response variables between female and male bees, we used a subset of our data containing only non-eusocial bees since all sampled bumblebees and honeybees were females. We also tested whether pollen species co-transport degree and strength, extracted from pollen co-transport networks, varied among pollinator groups using GLMM with negative binomial and Gamma error-distributions, respectively. For both models, we included plant species identity as a random effect. For all models, likelihood-ratio tests were used to evaluate the significance of the fixed effects when comparing the goodness of fit between the models and their respective null model. *Post-hoc* comparisons between pollinator categories were conducted with the *multcomp* package (Hothorn *et al*., 2008). All data analyses were conducted in R v.4.2.3 (R Core Team, 2023).

## 3 Results

### 3.1 Insect and pollen load diversity

In total, 781 flower-visiting insects were collected at both studied sites (RHA = 569, BS = 212) from which 710 were bees and 71 flies. Our samples represented 134 morphospecies, including 100 bee (29 genera) and 34 fly morphospecies (Table S1). Bee samples comprised 51 bumblebees (five *Bombus* spp., mainly *Bombus vosnesenskii*), 71 honeybees, 220 megachilid bees (10 genera and 48 morphospecies), and 368 other non-eusocial bees (17 genera and 46 morphospecies) (Table S1). We collected 475 females and 113 males, disregarding eusocial bees, which are represented by female foragers only. Overall, we counted 273,157 pollen grains from insect pollen loads representing 41 plant species (Table S2). Insects carried, on average, 349 pollen grains (median = 84) and a mean richness equivalent to three plant species (mean = 3.09, median = 3). The most frequent plant species observed in pollen load samples were *Antirrhinum cornutum* (Plantaginaceae) (36.3%), *Streptanthus breweri* (Brassicaceae) (32.6%) and *Mimulus guttatus* (Phrymaceae) (26.5%). Pure pollen loads (*i.e*., only containing a single plant species) represented 18% of samples, mainly containing pollen from *Clarkia gracilis* (Onagraceae), *Eriophyllum lanatum* (Asteraceae) and *S. breweri* (Table S2) and were found in several morphospecies of bees (36) and flies (10). Only 6.5% of insect samples contained no pollen grains. This percentage was also represented by different morphospecies of flower-visiting bees (25) and flies (8).

### 3.2 Variation in pollen transport components among pollinator groups

Pollinator groups differed in pollen load size (χ^2^ = 11.77, *df* = 4, *p* = 0.019; Figure 2A), pollen load richness (χ^2^ = 12.83, *df* = 4, *p* = 0.012; Figure 2B) and MPD (phylogenetic diversity) (χ^2^ = 14.10, *df* = 4, *p* = 0.007; Figure 2C), but not in MFTD (phenological diversity, range = 0.08-0.82) (χ^2^ = 4.50, *df* = 4, *p* = 0.343). Specifically, bumblebees significantly transported larger pollen loads than megachilid bees and flies, which in turn did not differ from honeybees and other bees (Figure 2A). No differences in pollen load size were detected among bumblebees, honeybees, and other bees (Figure 2A). A larger pollen load richness was observed for bumblebees compared to other bees and flies (Figure 2B). No pairwise differences in pollen load richness were found between honeybees, megachilid bees, other bees, and flies (Figure 2B). Pairwise comparisons between insect groups indicated a significantly higher MPD for megachilid bees compared to flies (Figure 2C). A similar but marginally significant pattern was observed for bumble bees compared to flies (Figure 2C). None other pairwise comparisons were significant.

**Figure 2.**
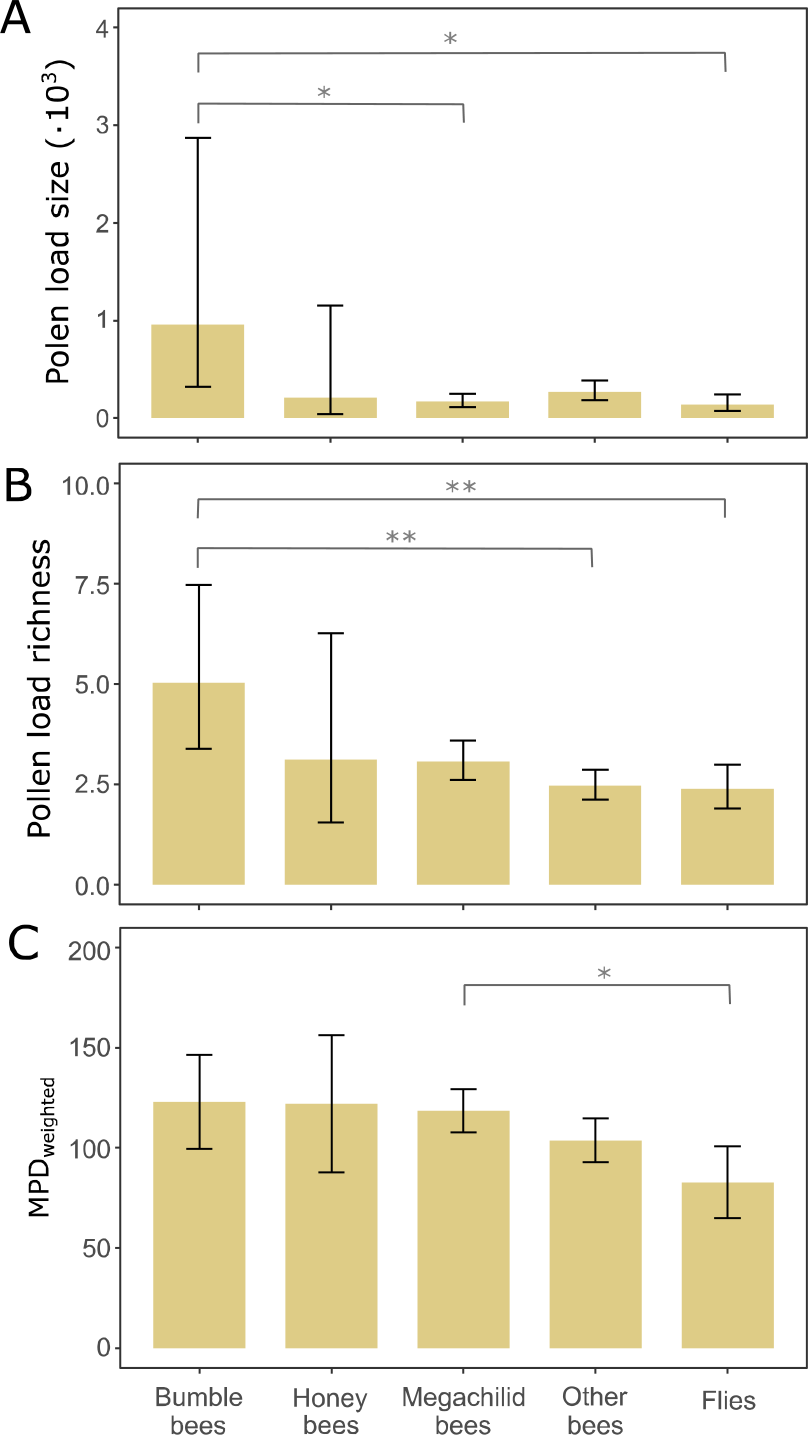
Least-squares means (±95% CI) of pollen load size (A), richness (B). and phylogenetical diversity (C), represented by the weighted mean phylogenetic distance (MPD_weighted_) obtained for each pollinator group in the co-flowering community at the serpentine seeps. Significance (*p*-value): < .001^***^, < .010^**^, < .050^*^.

### 3.3 Pollinator body size and sex effects on pollen transport

Bee body size was positively related to both, pollen load size (χ^2^ = 11.57, *df* = 4, *p* = 0.021; Figure 3A) and richness (χ^2^ = 15.15, *df* = 4, *p* = 0.004; Figure 3B), and these relationships did not vary with bee group (pollen load size: χ^2^ = 1.00, *df* = 3, *p* = 0.802; pollen load richness: χ^2^ = 4.36, *df* = 3, *p* = 0.225). Body size predicted neither MPD (χ^2^ = 1.25, *df* = 1, *p* = 0.263) nor MFTD (χ^2^ = 0.72, *df* = 1, *p* = 0.396).

**Figure 3.**
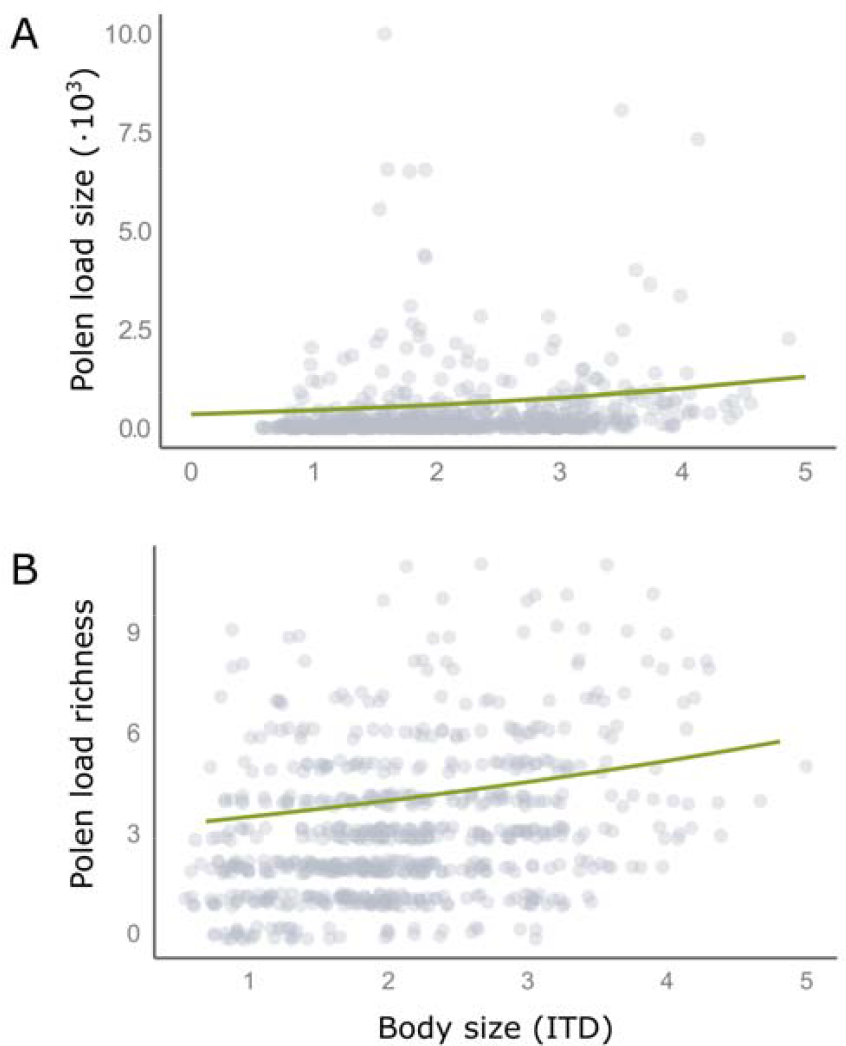
Effect of body size, measured as the intertegular distance of bees (ITD), on pollen load size (*P* < 0.001) and richness (*P* = 0.362) carried by bees at the serpentine seeps.

Disregarding eusocial bees and flies, sex was also an important driver of pollen load size (χ^2^ = 15.77, *df* = 1, *p* < 0.001; Figure 4A), but not pollen load richness (χ^2^ = 0.83, *df* = 1, *p* = 0.3). Female pollen load size was more than two times larger compared to that of males (Figure 4A), even though pollen load richness of both groups was similar. However, males carried pollen loads with higher MPD than females (χ^2^ = 4.17, *df* = 1, *p* = 0.041; Figure 4B), whilst MFTD was similar between sexes (χ^2^ = 0.23, *df* = 1, *p* = 0.631).

**Figure 4.**
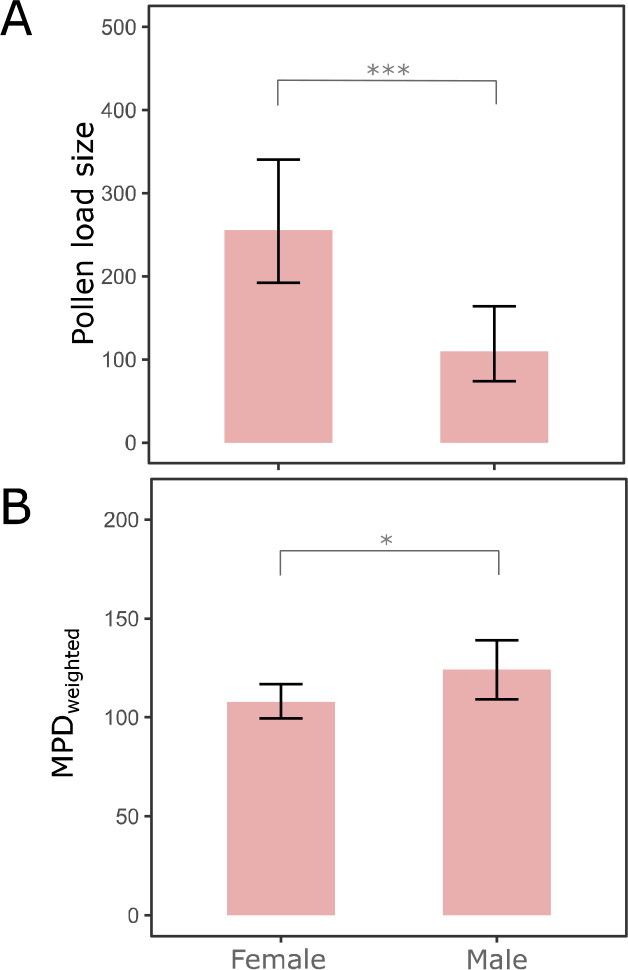
Least-squares means (±95% CI) of pollen load size (A) and phylogenetical diversity (B), represented by the weighted mean phylogenetic distance (MPD_weighted_) carried by female and male bees at the serpentine seeps. Significance (*p*-value): < .001^***^, < .010^**^, < .050^*^.

### 3.4 Pollen co-transport networks of pollinator groups

We constructed pollen co-transport networks for 281 flower-visiting individual insects in four pollinator groups (excluding other bees) and that were collected only at RHA (Figure 5). We observed a total of 38 pollen species that were co-transported (*i.e*., shared an insect body) with other species. Co-transport networks contained between 32-25 pollen species across pollinator groups (bumblebees = 29, honeybees = 30, megachilid bees = 32, and flies = 25) (Figure 5). The overall size of pollen co-transport networks (*i.e*., number of links) varied greatly among pollinator groups and ranged from 91 (flies) to 281 (megachilid bees). Network modularity also varied among pollinator groups ranging from 0.30 (megachilid bees) to 0.54 (flies) indicating qualitative differences in the number of co-transport network modules formed across pollinators groups (4-6 modules; Figure 5). Modules contained, on average, four to six pollen species per pollinator group (Figure 5).

**Figure 5.**
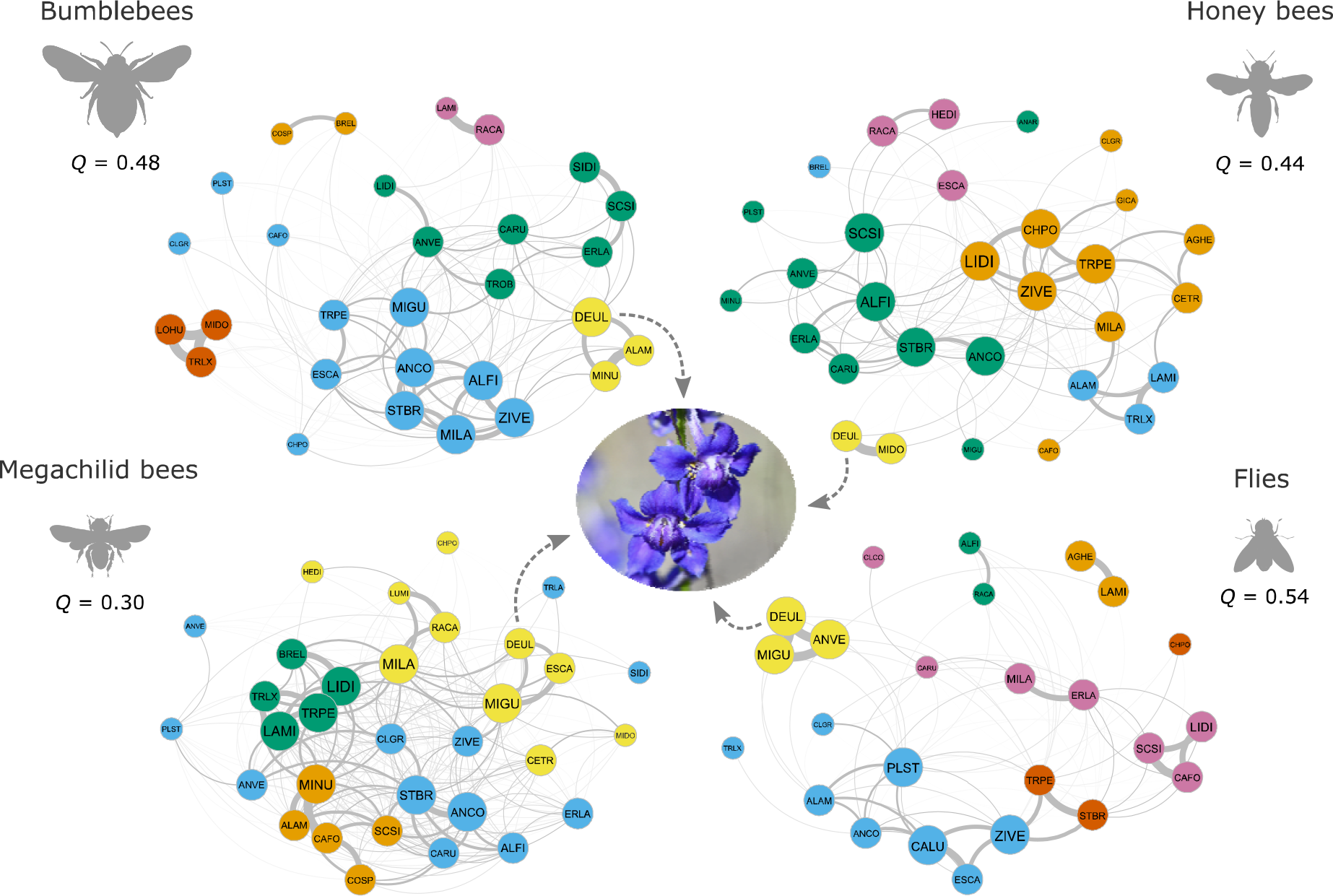
Pollen co-transport networks for four pollinator groups (bumblebees, honeybees, megachilid bees and flies). The modularity index (*Q*) for each network is shown. Each node represents a pollen species and its size represents the quantiles of pollen co-transport strength (weighted degree). Edge width represents the Schoener’s insect body use overlap index (co-transport index) between pollen species-pairs. The different colors represent different pollen co-transport modules within each pollinator group. Note that *Delphinium uliginosum* (DEUL shown at the center) travels with 2-9 other pollen species depending on the pollinator group. Species codes are note inside each node and full species names and codes are given in Table S2.

Pollinator groups differ significantly in their pollen co-transport degree (χ^2^ = 47.45, *df* = 3, *p* < 0.001; Figure 6), but not in co-transport strength (range = 0.04-3.58) (χ^2^ = 3.11, *df* = 3, *p* = 0.3). Megachilid bees had a significantly higher pollen co-transport degree, indicating that pollen carried by megachilid bees was co-transported with more species than pollen grains on bumblebees, honeybees, and flies (Figure 6). On the contrary, pollen species carried by flies and honeybees were co-transported with a lesser number of other pollen species (low co-transport degree) compared to other pollinator groups (Figure 6).

**Figure 6.**
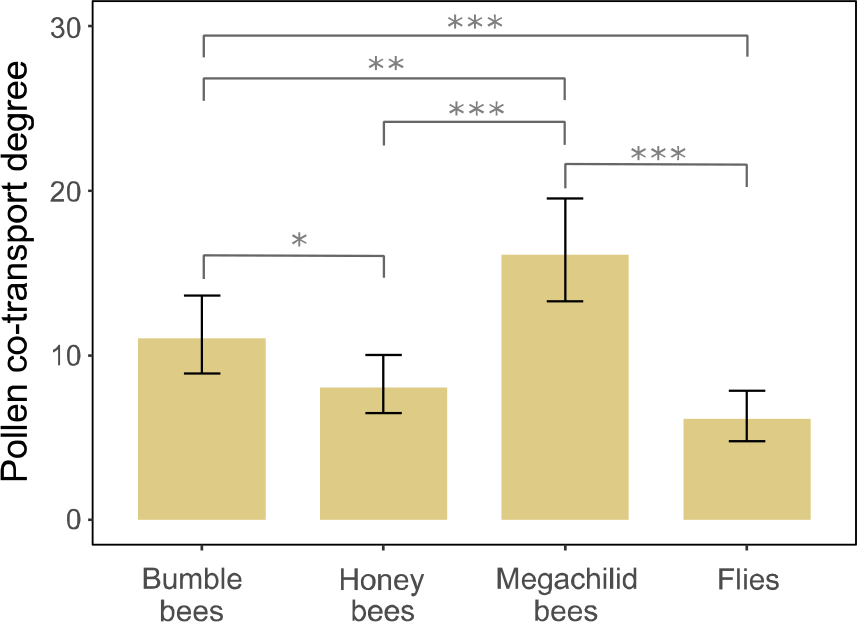
Least-squares means (±95% CI) of pollen co-transport degree that indicates the average number of co-transport partners of species per pollinator group in the co-flowering community. Significance (*p*-value): < .001^***^, < .010^**^, < .050^*^.

## 4 Discussion

Individual insects carried on average between 7-12% (∼3-5 species) of the total number of ‘pollen species’ (41 species) transported by the insect groups studied, suggesting a more narrow niche breadth at the individual level (*e.g*., Smith *et al*., 2019). We also observed differences in patterns of pollen transport (load size, richness, MPD) and co-transport (co-transport network degree and modularity) among pollinator groups, and hence in the potential for pollen–pollen competitive interactions on insect bodies within (inter-individual) and between pollinator groups. The ‘competitive landscape’ that pollen species face during transport is further shaped by morphological, ecological and behavioral characteristics intrinsic to different pollinator groups. Specifically, taxonomic identity, body size and sex of insects can influence the pollen ‘landscape’ on their bodies (pollen load size, richness, and phylogenetic diversity). Our results provide evidence suggesting that different pollinator groups represent vastly different ‘pollen competitive arenas’, that may impose a different set of ecological and evolutionary pressures on pollen species as they travel on insect bodies, with unknown consequences for plant species and plant community assembly.

At our study sites, bumblebees carried the largest and most diverse pollen loads. However, we also observed large within-group variation in pollen load size and richness in bumblebees. This large within-group variation may suggest differences in individual foraging that are not captured when evaluating group/species-level patterns, but that could play an important role in the structure of community-level plant–pollinator interactions (Olesen *et al*., 2010; Tur *et al*., 2014; Brosi, 2016). In fact, pollen co-transport networks revealed high modularity within bumblebees (six modules) and a large variation in the number of co-transported pollen species within each module (2-12 pollen species), suggesting a substructure of pollen–pollen interactions at the individual level within this group (Figure 5). Nonetheless, significant differences in pollen load size were observed between bumblebees, megachilid bees, and flies. We also observed differences in pollen load richness between bumblebees, flies and other bees, hence supporting their high relevance as pollinators (Memmott *et al*., 2004; Goulson, 2006). The high diversity in the pollen loads carried by bumblebees, however, could also help strengthen plant–plant competitive interactions via heterospecific pollen transfer (Morales & Traveset, 2008; Cullen *et al*., 2021), with detrimental effects for plant fitness (Morales & Traveset, 2008; Moreira-Hernández & Muchhala, 2019; Arceo-Gómez, 2021). Thus, either via their positive or negative effects, bumblebees may play a large role in co-flowering community assembly at the study sites.

Interestingly, the pollen load size and richness carried by honeybees was equivalent to that of any other pollinator group, revealing similar roles in pollen transport and suggesting that honeybees may not be more important than other insect groups in their role as pollinators (Geslin *et al*., 2017; Magrach *et al*., 2017; Travis & Kohn, 2023). On the contrary, our results show that understudied groups such as megachilid bees and flies may play an underappreciated role as pollinators, as they carry comparable pollen loads in terms of size and diversity to other insect groups. Specifically, flies transport pollen loads similar in size and richness as non-bumblebee groups, and these are as phylogenetically diverse as those of bumblebees and honeybees. Flies, however, carry less phylogenetically diverse pollen loads compared to megachilid bees, suggesting differences in the evolutionary history of the plants they visit. Other studies in the same community have found similar results (Cullen *et al*., 2021), suggesting that flies may be important pollinators in this, and perhaps other highly diverse co-flowering communities.

Pollen co-transport networks revealed that the number (*i.e*., network degree) and identity of potential competitors that individual pollen species encounter on insect bodies varies across pollinator groups. Pollinator groups hence likely represent different ‘competitive arenas’ for pollen species. For instance, pollen species traveling on bodies of megachilid bees can encounter and compete with almost twice the number of pollen species (15), compared to those traveling on flies (8; Figure 6). Our co-transport network approach also showed that megachilid bees generate the highest diversity of pollen–pollen interactions, even though bumblebees transport the largest and most diverse pollen loads. This may result from differences in foraging niche partitioning among individuals within each group, or to the high diversity of megachilid bees in the community. Nevertheless, it suggests that a high pollen load richness at the species or group level may not directly translate to a higher diversity of realized pollen–pollen interactions on insect bodies (*e.g*., Cullen *et al*. 2021) when we consider patterns of pollen transport at the individual level. Furthermore, our network approach revealed that some pollen species travel, and thus likely compete more often with each other on a pollinator’s body (*i.e*., co-transport modules) (Figure 5), but that a single pollen species may face different ‘competitive arenas’ (*i.e*., number and identity of pollen competitors) as it is transported by different pollinator groups. For example, *Delphinium uliginosum* (Ranuculaceae) pollen travels consistently with only one other pollen species (*Minuartia douglasii*; Caryophyllaceae) when transported by honeybees, or with nine others when transported by megachilid bees (Figure 5), thus facing very different competitive environments. Overall, our results suggest that differences in pollinator attributes and behaviors can lead to differences in the opportunity for plant species to compete during pollen co-transport. These differences in the pollen competition landscape on insect bodies (Minnaar *et al*., 2019) within and among different pollinator groups may have unforeseen ecological and evolutionary consequences. For instance, stronger competition for space on pollinator bodies could lead to the evolution of pollen weaponry (*e.g*., ornaments and chemicals) (Murphy, 2000; Minnaar *et al*., 2019), divergence on pollen placement (*e.g*., diffuse, stroke, stamp or layered) (Morris *et al*., 1995; Minnaar *et al*., 2019; Moir & Anderson, 2023) and/or dispersal strategies (*e.g*., sequential or vertical) (Harder & Wilson, 1998; Minnaar *et al*., 2019) that potentially maximize siring success. If the consequences for male (*e.g*., Moreira-Hernández and Muchhala 2019) and female fitness (*e.g*., Arceo-Gómez & Ashman, 2011) in plants are strong, it may also lead to divergence or shifts in pollinator assemblages. Thus, evaluating the full complexity of pollen ‘landscapes’ on insect bodies within and among a diversity of pollinator groups is central for understanding their importance not only as pollinators, but also as mediators of competitive plant–plant interactions via male (pollen competition on insect bodies) and female fitness (heterospecific pollen deposition), with potential consequences for plant coexistence and community assembly.

Our results also revealed that pollen load characteristics can vary according to pollinator body size and sex-related differences. Pollinator body size has been widely recognized as a trait affecting foraging range (Greenleaf *et al*., 2007; Wright *et al*., 2015), floral fit (Agosta & Janzen, 2005; Solís-Montero & Vallejo-Marín, 2017), pollen deposition on stigmas (Földesi *et al*., 2021) and pollination effectiveness (Jauker *et al*., 2016). Here, we observed a positive relationship between body size and pollen load size and richness also found by previous studies (Smith *et al*., 2019; Cullen *et al*., 2021). These results combined suggest that the importance of pollinators for plant reproductive success and as mediators of plant–plant interactions is dependent on body size. This is key as recent studies have found steep decreases in pollinator body size as a result of increasing temperatures (Herrera *et al*., 2023) and habitat disturbance (Grab et al. 2019, Fitzgerald et al. 2022; but see Warzecha et al. 2016,). So far, the effects of warmer temperatures and habitat fragmentation on pollination have been mainly attributed to changes in phenology leading to plant–pollinator mismatches (Forrest, 2015) and to pollinator declines (Klein *et al*., 2007; Potts *et al*., 2010). However, it is possible that warmer temperatures and habitat disturbances will also affect patterns of pollen transport and co-transport, pollinator efficiency and pollinator niche breadth via changes in insect body size, with unexplored consequences for the long-term persistence of plant communities.

Female bees carried twice the amount of pollen than males which may be explained by between-sex differences in foraging behavior and ecological role (Ne’eman *et al*., 2006; Roswell *et al*., 2019). Whilst females show a more prominent and active role in pollen collection (Tang *et al*., 2019), males use flowers to feed on nectar, mate search and rest (Eickwort & Ginsberg, 1980; Pinheiro *et al*., 2017; Danforth, 2019). However, we found no differences in pollen load richness carried between male and females, suggesting a similar niche breadth size for both sexes. It is important to note that in this study we avoided collecting pollen from pollen-carrying structures (*i.e*., corbiculae or scopae) and thus pollen on insect bodies represents pollen that is relevant for pollination. Thus, sex-specific differences in pollen transport (load size) may have consequences for pollinator-mediated plant–plant interactions (Nakazawa & Kishi, 2023). Interestingly, pollen loads of males were more phylogenetically diverse than those of females despite similar pollen species richness, indicating differences in the evolutionary history and composition of the plants they visit (Roswell *et al*., 2019). Specifically, females visit a more closely set of related plants compared to male bees (also see Cullen *et al*., 2021), which can be to some extent a result of floral constancy or oligolectic levels that narrow the foraging for pollen to related species in many solitary female bees (Schlindwein, 2004; Cane & Sipes, 2006). In contrast, males carrying pollen of more distantly related species may be explained by the diversity of male activities on flowers other than foraging (Danforth, 2019). This result also suggests that female-mediated heterospecific pollen transfer may have stronger detrimental consequences for plant fitness compared to male-mediated transfer. Negative fitness effects of heterospecific pollen transfer have been shown to be stronger between closely related species (ArceolGómez & Ashman, 2016; Streher *et al*., 2020). However, to our knowledge, differential effects of heterospecific pollen transfer driven by male and female bees in natural communities is a tantalizing possibility that remains unexplored.

Finally, we did not observe differences in the phenological diversity (flowering time distance) of pollen loads due to pollinator group, body size or sex suggesting no differences in the temporal niche breadth of pollinators despite differences in life history strategies among groups (*e.g*., eusocial vs non-eusocial). The absence of differences may be the result of a relatively short flowering season (∼2-3 months). What is evident, however, is the importance of evaluating patterns of pollen transport beyond just pollen diversity and abundance. And that pollen co-transport networks can provide key insights that help inform on the variation in the identity and intensity of pollen–pollen interactions within and across pollinator groups. Overall, our findings emphasize the importance of evaluating factors that affect intermediate stages in the pollination process, which are often overlooked and have the potential to inform on key ecological and evolutionary process within communities.

## Data availability

The dataset used in this study will be deposited in an open digital repository (Dryad) upon acceptance.

## Acknowledgements

This work was supported by the National Science Foundation (DEB 1931163 to GAG). We thank K. Riddle, J. Horton, J. Bailey and M. Merkel for their help collecting data in the field and in the lab.

## Conflict of interest

The authors have no conflicts of interest to declare that are relevant to the content of this article.

## Author contributions

LTC and GAG conceived the study; JNW, DAB and GAG conducted fieldwork; LTC, JNW, CM and JWA conducted lab work; LTC led the analyses; and LTC and GAG wrote the manuscript with substantial contributions by all other authors. All authors gave final approval for publication.

## Notes

### Competing Interest Statement

The authors have declared no competing interest.

