## Supporting Information for "Patterns and drivers of pollen co-transport networks vary across pollinator groups"

**Table S1.** List of flower-visiting morphospecies of bees and flies and number of individuals per morphospecies sampled at the studied serpentine seep plant meta-community in Northern California, USA, over 13 days between May 9th and June 1st, during the peak flowering season in 2021.

| **Insect order** | **Family** | **Genus** | **Morphospecies/species** | **Pollinator group** | **Number of individuals** |
| --- | --- | --- | --- | --- | --- |
| Hymenoptera | Andrenidae | *Andrena* | *Andrena astragali* | Other bees | 23 |
| Hymenoptera | Andrenidae | *Andrena* | *Andrena subchalybea* | Other bees | 77 |
| Hymenoptera | Andrenidae | *Andrena* | *Andrena* sp.1 | Other bees | 2 |
| Hymenoptera | Andrenidae | *Andrena* | *Andrena* sp.2 | Other bees | 2 |
| Hymenoptera | Andrenidae | *Andrena* | *Andrena* sp.3 | Other bees | 8 |
| Hymenoptera | Andrenidae | *Calliopsis* | *Calliopsis* sp.1 | Other bees | 9 |
| Hymenoptera | Andrenidae | *Calliopsis* | *Calliopsis* sp.2 | Other bees | 1 |
| Hymenoptera | Apidae | *Anthophora* | *Anthophora* sp.1 | Other bees | 1 |
| Hymenoptera | Apidae | *Anthophora* | *Anthophora* sp.2 | Other bees | 2 |
| Hymenoptera | Apidae | *Anthophora* | *Anthophora* sp.3 | Other bees | 1 |
| Hymenoptera | Apidae | *Apis* | *Apis mellifera* | Honey bees | 71 |
| Hymenoptera | Apidae | *Bombus* | *Bombus vosnesenskii* | Bumblebees | 38 |
| Hymenoptera | Apidae | *Bombus* | *Bombus* sp.1 | Bumblebees | 9 |
| Hymenoptera | Apidae | *Bombus* | *Bombus* sp.2 | Bumblebees | 2 |
| Hymenoptera | Apidae | *Bombus* | *Bombus* sp.3 | Bumblebees | 1 |
| Hymenoptera | Apidae | *Bombus* | *Bombus* sp.4 | Bumblebees | 1 |
| Hymenoptera | Apidae | *Ceratina* | *Ceratina* sp.1 | Other bees | 3 |
| Hymenoptera | Apidae | *Ceratina* | *Ceratina* sp.2 | Other bees | 3 |
| Hymenoptera | Apidae | *Diadasia* | *Diadasia bituberculata* | Other bees | 4 |
| Hymenoptera | Apidae | *Diadasia* | *Diadasia nigrifrons* | Other bees | 1 |
| Hymenoptera | Apidae | *Diadasia* | *Diadasia* sp.1 | Other bees | 1 |
| Hymenoptera | Apidae | *Diadasia* | *Diadasia* sp.2 | Other bees | 22 |
| Hymenoptera | Apidae | *Diadasia* | *Diadasia* sp.3 | Other bees | 2 |
| Hymenoptera | Apidae | *Diadasia* | *Diadasia* sp.4 | Other bees | 4 |
| Hymenoptera | Apidae | *Diadasia* | *Diadasia* sp.5 | Other bees | 1 |
| Hymenoptera | Apidae | *Epeoloides* | *Epeoloides* sp.1 | Other bees | 1 |
| Hymenoptera | Apidae | *Eucera* | *Eucera* sp.1 | Other bees | 5 |
| Hymenoptera | Apidae | *Eucera* | *Eucera* sp.2 | Other bees | 16 |
| Hymenoptera | Apidae | *Habropoda* | *Habropoda* sp.1 | Other bees | 1 |
| Hymenoptera | Apidae | *Habropoda* | *Habropoda* sp.2 | Other bees | 1 |
| Hymenoptera | Apidae | *Melissodes* | *Melissodes lupina* | Other bees | 1 |
| Hymenoptera | Apidae | *Melissodes* | *Melissodes lustra* | Other bees | 14 |
| Hymenoptera | Apidae | *Nomada* | *Nomada* sp.1 | Other bees | 1 |
| Hymenoptera | Apidae | *Nomada* | *Nomada* sp.2 | Other bees | 1 |
| Hymenoptera | Apidae | *Nomada* | *Nomada* sp.3 | Other bees | 1 |
| Hymenoptera | Apidae | *Triepeolus* | *Triepeolus* sp.1 | Other bees | 2 |
| Hymenoptera | Apidae | *Xeromelecta* | *Xeromelecta* sp.1 | Other bees | 1 |
| Hymenoptera | Apidae | *Xylocopa* | *Xylocopa* sp.1 | Other bees | 2 |
| Hymenoptera | Colletidae | *Hylaeus* | *Hylaeus* sp.1 | Other bees | 1 |
| Hymenoptera | Halictidae | *Dufourea* | *Dufourea* sp.1 | Other bees | 2 |
| Hymenoptera | Halictidae | *Halictus* | *Halictus* sp.1 | Other bees | 2 |
| Hymenoptera | Halictidae | *Halictus* | *Halictus* sp.2 | Other bees | 8 |
| Hymenoptera | Halictidae | *Halictus* | *Halictus* sp.3 | Other bees | 4 |
| Hymenoptera | Halictidae | *Halictus* | *Halictus* sp.4 | Other bees | 1 |
| Hymenoptera | Halictidae | *Halictus* | *Halictus* sp.5 | Other bees | 42 |
| Hymenoptera | Halictidae | *Lasioglossum* | *Lasioglossum* sp.1 | Other bees | 11 |
| Hymenoptera | Halictidae | *Lasioglossum* | *Lasioglossum* sp.2 | Other bees | 1 |
| Hymenoptera | Halictidae | *Lasioglossum* | *Lasioglossum* sp.3 | Other bees | 1 |
| Hymenoptera | Halictidae | *Lasioglossum* | *Lasioglossum* sp.4 | Other bees | 2 |
| Hymenoptera | Halictidae | *Lasioglossum* | *Lasioglossum* sp.5 | Other bees | 75 |
| Hymenoptera | Halictidae | *Lasioglossum* | *Lasioglossum* sp.6 | Other bees | 1 |
| Hymenoptera | Halictidae | *Lasioglossum* | *Lasioglossum* sp.7 | Other bees | 4 |
| Hymenoptera | Megachilidae | *Anthidiellum* | *Anthidiellum* sp.1 | Megachilid bees | 8 |
| Hymenoptera | Megachilidae | *Anthidium* | *Anthidium* sp.1 | Megachilid bees | 2 |
| Hymenoptera | Megachilidae | *Anthidium* | *Anthidium* sp.2 | Megachilid bees | 4 |
| Hymenoptera | Megachilidae | *Anthidium* | *Anthidium* sp.3 | Megachilid bees | 3 |
| Hymenoptera | Megachilidae | *Anthidium* | *Anthidium* sp.4 | Megachilid bees | 2 |
| Hymenoptera | Megachilidae | *Anthidium* | *Anthidium* sp.5 | Megachilid bees | 19 |
| Hymenoptera | Megachilidae | *Ashmeadiella* | *Ashmeadiella* sp.1 | Megachilid bees | 14 |
| Hymenoptera | Megachilidae | *Ashmeadiella* | *Ashmeadiella* sp.2 | Megachilid bees | 1 |
| Hymenoptera | Megachilidae | *Ashmeadiella* | *Ashmeadiella* sp.3 | Megachilid bees | 1 |
| Hymenoptera | Megachilidae | *Ashmeadiella* | *Ashmeadiella* sp.4 | Megachilid bees | 1 |
| Hymenoptera | Megachilidae | *Ashmeadiella* | *Ashmeadiella* sp.5 | Megachilid bees | 1 |
| Hymenoptera | Megachilidae | *Ashmeadiella* | *Ashmeadiella* sp.6 | Megachilid bees | 1 |
| Hymenoptera | Megachilidae | *Ashmeadiella* | *Ashmeadiella* sp.7 | Megachilid bees | 1 |
| Hymenoptera | Megachilidae | *Ashmeadiella* | *Ashmeadiella* sp.8 | Megachilid bees | 2 |
| Hymenoptera | Megachilidae | *Ashmeadiella* | *Ashmeadiella* sp.9 | Megachilid bees | 2 |
| Hymenoptera | Megachilidae | *Chelostoma* | *Chelostoma* sp.1 | Megachilid bees | 1 |
| Hymenoptera | Megachilidae | *Chelostoma* | *Chelostoma* sp.2 | Megachilid bees | 1 |
| Hymenoptera | Megachilidae | *Chelostoma* | *Chelostoma* sp.3 | Megachilid bees | 1 |
| Hymenoptera | Megachilidae | *Coelioxys* | *Coelioxys* sp.1 | Megachilid bees | 2 |
| Hymenoptera | Megachilidae | *Coelioxys* | *Coelioxys* sp.2 | Megachilid bees | 1 |
| Hymenoptera | Megachilidae | *Dianthidium* | *Dianthidium* sp.1 | Megachilid bees | 25 |
| Hymenoptera | Megachilidae | *Hoplitis* | *Hoplitis* sp.1 | Megachilid bees | 1 |
| Hymenoptera | Megachilidae | *Hoplitis* | *Hoplitis* sp.2 | Megachilid bees | 1 |
| Hymenoptera | Megachilidae | *Hoplitis* | *Hoplitis* sp.3 | Megachilid bees | 2 |
| Hymenoptera | Megachilidae | *Hoplitis* | *Hoplitis* sp.4 | Megachilid bees | 6 |
| Hymenoptera | Megachilidae | *Hoplitis* | *Hoplitis* sp.5 | Megachilid bees | 1 |
| Hymenoptera | Megachilidae | *Hoplitis* | *Hoplitis* sp.6 | Megachilid bees | 1 |
| Hymenoptera | Megachilidae | *Hoplitis* | *Hoplitis* sp.7 | Megachilid bees | 1 |
| Hymenoptera | Megachilidae | *Hoplitis* | *Hoplitis* sp.8 | Megachilid bees | 2 |
| Hymenoptera | Megachilidae | *Hoplitis* | *Hoplitis* sp.9 | Megachilid bees | 1 |
| Hymenoptera | Megachilidae | *Hoplitis* | *Hoplitis* sp.10 | Megachilid bees | 5 |
| Hymenoptera | Megachilidae | *Hoplitis* | *Hoplitis* sp.11 | Megachilid bees | 1 |
| Hymenoptera | Megachilidae | *Megachile* | *Megachile* sp.1 | Megachilid bees | 5 |
| Hymenoptera | Megachilidae | *Megachile* | *Megachile* sp.2 | Megachilid bees | 2 |
| Hymenoptera | Megachilidae | *Megachile* | *Megachile* sp.3 | Megachilid bees | 2 |
| Hymenoptera | Megachilidae | *Megachile* | *Megachile* sp.4 | Megachilid bees | 1 |
| Hymenoptera | Megachilidae | *Megachile* | *Megachile* sp.5 | Megachilid bees | 1 |
| Hymenoptera | Megachilidae | *Megachile* | *Megachile* sp.6 | Megachilid bees | 1 |
| Hymenoptera | Megachilidae | *Megachile* | *Megachile* sp.7 | Megachilid bees | 2 |
| Hymenoptera | Megachilidae | *Megachile* | *Megachile* sp.8 | Megachilid bees | 3 |
| Hymenoptera | Megachilidae | *Osmia* | *Osmia* sp.1 | Megachilid bees | 16 |
| Hymenoptera | Megachilidae | *Osmia* | *Osmia* sp.2 | Megachilid bees | 31 |
| Hymenoptera | Megachilidae | *Osmia* | *Osmia* sp.3 | Megachilid bees | 22 |
| Hymenoptera | Megachilidae | *Osmia* | *Osmia* sp.4 | Megachilid bees | 3 |
| Hymenoptera | Megachilidae | *Osmia* | *Osmia* sp.5 | Megachilid bees | 4 |
| Hymenoptera | Megachilidae | *Osmia* | *Osmia* sp.6 | Megachilid bees | 4 |
| Hymenoptera | Megachilidae | *Osmia* | *Osmia* sp.7 | Megachilid bees | 6 |
| Hymenoptera | Megachilidae | *Protosmia* | *Protosmia* sp.1 | Megachilid bees | 1 |
| Diptera | - | - | Diptera sp.1 | Flies | 1 |
| Diptera | - | - | Diptera sp.2 | Flies | 5 |
| Diptera | - | - | Diptera sp.3 | Flies | 2 |
| Diptera | - | - | Diptera sp.4 | Flies | 1 |
| Diptera | - | - | Diptera sp.5 | Flies | 1 |
| Diptera | - | - | Diptera sp.6 | Flies | 1 |
| Diptera | - | - | Diptera sp.7 | Flies | 3 |
| Diptera | - | - | Diptera sp.8 | Flies | 1 |
| Diptera | - | - | Diptera sp.9 | Flies | 1 |
| Diptera | - | - | Diptera sp.10 | Flies | 1 |
| Diptera | - | - | Diptera sp.11 | Flies | 1 |
| Diptera | - | - | Diptera sp.12 | Flies | 1 |
| Diptera | - | - | Diptera sp.13 | Flies | 2 |
| Diptera | - | - | Diptera sp.14 | Flies | 1 |
| Diptera | - | - | Diptera sp.15 | Flies | 2 |
| Diptera | - | - | Diptera sp.16 | Flies | 1 |
| Diptera | - | - | Diptera sp.17 | Flies | 1 |
| Diptera | - | - | Diptera sp.18 | Flies | 1 |
| Diptera | - | - | Diptera sp.19 | Flies | 2 |
| Diptera | - | - | Diptera sp.20 | Flies | 4 |
| Diptera | - | - | Diptera sp.21 | Flies | 1 |
| Diptera | - | - | Diptera sp.22 | Flies | 1 |
| Diptera | - | - | Diptera sp.23 | Flies | 1 |
| Diptera | - | - | Diptera sp.24 | Flies | 1 |
| Diptera | - | - | Diptera sp.25 | Flies | 1 |
| Diptera | - | - | Diptera sp.26 | Flies | 1 |
| Diptera | - | - | Diptera sp.27 | Flies | 1 |
| Diptera | - | - | Diptera sp.28 | Flies | 1 |
| Diptera | - | - | Diptera sp.29 | Flies | 1 |
| Diptera | - | - | Diptera sp.30 | Flies | 22 |
| Diptera | - | - | Diptera sp.31 | Flies | 1 |
| Diptera | - | - | Diptera sp.32 | Flies | 4 |
| Diptera | - | - | Diptera sp.33 | Flies | 1 |
| Diptera | - | - | Diptera sp.34 | Flies | 1 |
|  |  |  |  | **Total** | **781** |

**Table S2.** List of plant species found within pollen loads of the sampled flower-visiting insects at the studied serpentine seep plant meta-community in Northern California, USA.

| **Plant Species Code** | **Plant Species** | **Plant Family** |
| --- | --- | --- |
| ACMI | *Achillea millefolium* | Asteraceae |
| AGHE | *Agoseris heterophylla* | Asteraceae |
| ALAM | *Allium amplectens* | Amaryllidaceae |
| ALFI | *Allium fimbriatum* | Amaryllidaceae |
| ANAR | *Anagallis arvensis* | Primulaceae |
| ANCO | *Antirrhinum cornutum* | Plantaginaceae |
| ANVE | *Antirrhinum vexillocalyculatum* | Plantaginaceae |
| BREL | *Brodiaea elegans* | Asparagaceae |
| CAAM | *Calochortus amabilis* | Liliaceae |
| CAFO | *Castilleja foliolosa* | Orobanchaceae |
| CALU | *Calochortus luteus* | Liliaceae |
| CARU | *Castilleja rubicundula* | Orobanchaceae |
| CETR | *Centaurium trichanthum* | Gentianaceae |
| CHPO | *Chlorogalum pomeridianum* | Asparagaceae |
| CLCO | *Clarkia concinna* | Onagraceae |
| CLGR | *Clarkia gracilis* | Onagraceae |
| COSP | *Collinsia sparsiflora* | Plantaginaceae |
| DEUL | *Delphinium uliginosum* | Ranunculaceae |
| DIVO | *Dichelostemma volubile* | Asparagaceae |
| ERLA | *Eriophyllum lanatum* | Asteraceae |
| ESCA | *Eschscholzia californica* | Papaveraceae |
| GICA | *Gilia capitata* | Polemoniaceae |
| HEDI | *Hesperolinon disjunctum* | Linaceae |
| LAMI | *Lagophylla minor* | Asteraceae |
| LIDI | *Linanthus dichotomus* | Polemoniaceae |
| LOHU | *Lotus humistratus* | Fabaceae |
| LUMI | *Lupinus microcarpus* | Fabaceae |
| MIGU | *Mimulus guttatus* | Phrymaceae |
| MILA | *Mimulus layneae* | Phrymaceae |
| MIDO | *Minuartia douglasii* | Caryophyllaceae |
| MINU | *Mimulus nudatus* | Phrymaceae |
| PLST | *Plagiobothrys stipitatus* | Boraginaceae |
| RACA | *Ranunculus californicus* | Ranunculaceae |
| SCSI | *Scutellaria siphocampyloides* | Lamiaceae |
| SIDI | *Sidalcea diploscypha* | Malvaceae |
| STBR | *Streptanthus breweri* | Brassicaceae |
| TRLA | *Triteleia laxa* | Asparagaceae |
| TRLX | *Trichostema laxum* | Lamiaceae |
| TROB | *Trifolium obtusiflorum* | Fabaceae |
| TRPE | *Triteleia peduncularis* | Asparagaceae |
| ZIVE | *Zigadenus venenosus* | Melanthiaceae |
